# Dynamic T-cell receptor repertoire remodeling is associated with murine and human lung allograft dysfunction

**DOI:** 10.64898/2026.11.20.768159

**Authors:** Yiqiao Wang, Vyacheslav Palchevskiy, Shahrzad Lari, Scott Palmer, Jamie L. Todd, Samuel S. Weigt, John A. Belperio

## Abstract

Chronic lung allograft dysfunction (CLAD) is a major cause of late graft failure after lung transplantation. Acute cellular rejection is an established risk factor for CLAD. However, its diagnosis by biopsy may incompletely capture dynamic and spatially heterogeneous immune remodeling within the allograft. We investigated whether lung graft proximal T-cell receptor beta-chain (TCRβ) repertoire features reflect alloimmune remodeling and associate with acute cellular rejection (ACR) and CLAD. TCRβ sequencing was performed on graft-infiltrating T-cells from a murine orthotopic lung transplant model and on 90 longitudinal bronchoalveolar lavage cell-pellet (BAL-cp) samples from 23 lung transplant recipients during the first post-transplant year. In the murine model, allografts showed persistent histologic injury and strong, consistent TCRβ repertoire remodeling relative to isografts. Across days 28 and 56, murine allografts had higher productive clonality (Cliff’s δ = 0.93, p = 0.0087) and maximum productive frequency (Cliff’s δ = 0.93, p = 0.014), with complete rank separation at each time point. Murine allografts also exhibited greater pairwise Morisita repertoire overlap (p < 0.0001) and enrichment of recurrent, high-frequency shared clonotypes. Productive clonality from human lung transplant recipients’ BAL-cp was higher at the a priori selected month-6 visit from patients who developed CLAD, as compared to those who remained CLAD-free (p = 0.0326). In a month-6 surveillance-visit index analysis, high productive clonality was associated with early onset CLAD (CLAD within 2 years of transplant) (log-rank p = 0.038). In a patient-specific index-event analysis, CLAD-free survival differed across high and low productive clonality groups among acute rejectors (log-rank p = 0.019). Together, these findings establish measurable graft-localized TCRβ repertoire remodeling and position BAL TCRβ profiling as a potential complement to biopsy diagnosis. With validation in larger cohorts, graft-proximal repertoire measures may help identify CLAD-associated immune states before the onset of lung function decline and inform more individualized surveillance.

## Introduction

Lung transplantation is often the final treatment option for end-stage lung diseases, but long-term survival is limited by chronic lung allograft dysfunction (CLAD), the leading cause of late graft failure after transplantation [1,2]. Although CLAD is multifactorial, T-cell–mediated alloimmunity is an important contributor to lung allograft injury [3–6]. Recipient T-cells recognize donor alloantigens, leading to clonal expansion and effector differentiation [6]. These responses also underlie acute cellular rejection (ACR), the T-cell–mediated form of acute allograft rejection [6,7]. Importantly, ACR is an established risk factor for subsequent CLAD, motivating investigation of the T-cell responses associated with acute rejection and chronic graft dysfunction [1,2,8].

ACR is diagnosed by transbronchial biopsy using the International Society for Heart and Lung Transplantation (ISHLT) histopathologic grading framework [7]. While essential for clinical diagnosis, biopsies survey a small portion of the allograft and are subject to sampling error and interobserver variability [9,10]. Moreover, histopathology provides a localized assessment of inflammation at a specific time point, but does not capture the clonal composition or longitudinal evolution of the underlying T-cell response. Graft-proximal repertoire profiling may therefore provide complementary biological information about immune repertoire dynamics that is not captured by histologic assessment alone. Sequencing the variable complementarity-determining region 3 (CDR3) of the T-cell receptor β chain (TCRβ) enables tracking of clonotypes across samples [11,12]. Repertoire analysis quantifies diversity, clonal expansion, dominant-clonotype frequency, and overlap between samples [11,12]. Productive clonality summarizes repertoire unevenness, with higher values indicating dominance by fewer clonotypes [13]. These measurements offer a means of monitoring antigen-driven T-cell responses after transplantation [13–16]. However, increased clonality is not specific for donor reactivity and may also reflect infection, inflammation, or other antigenic exposures [17]. TCR repertoire profiling is often performed on peripheral blood mononuclear cells (PBMCs), which may incompletely represent immune responses within the lung allograft. During ACR, expanded clonotypes detected in bronchoalveolar lavage (BAL) were discordant with circulating expansions and persisted as tissue-resident memory T-cells after treatment [18]. Longitudinal profiling also linked enrichment of host-versus-graft clones in BAL to rejection and pulmonary dysfunction [17]. In that study, graft–blood repertoire overlap was driven largely by nonalloreactive recipient CD8+ T-cells, further distinguishing circulating from intragraft responses [17]. Experimental evidence that alloreactive T-cells can be primed directly within the lung allograft further supports the importance of local immune responses [19]. BAL thus provides a clinically accessible window into graft-proximal T-cell responses, supporting longitudinal sampling in human recipients [3–5,20]. However, the relationship between BAL-derived productive clonality and subsequent CLAD, including in recipients with biopsy-proven ACR, remains incompletely defined.

In this study, we combined a murine orthotopic lung transplant model with an analysis of BAL cell pellet (BAL-cp) samples from human lung transplant recipients to investigate whether graft-proximal TCRβ repertoire remodeling reflects alloimmune activation and associates with ACR and CLAD. In mice, we compared allografts, isografts, and naïve lungs, hypothesizing that alloimmune injury would increase productive clonality, dominant-clonotype expansion, and repertoire convergence. Human analyses examined clonality during surveillance and at incident ACR to assess its association with subsequent CLAD.

## Materials and Methods

### Animals

Wild-type male BALB/c (H-2Kd) and C57BL/6 (H-2Kb) mice, 8–10 weeks old, were purchased from The Jackson Laboratories (Bar Harbor, ME, USA). Animals were housed under specific pathogen-free conditions in the UCLA animal facilities with access to food and water ad libitum.

### Orthotopic lung transplantation

Orthotopic left lung transplantation was performed by the cuff technique as described previously [21]. Briefly, C57BL/6 mice served as donors for isografts and BALB/c mice served as donors for allografts. Donor mice were anesthetized, intubated, mechanically ventilated, and the left lung was harvested. Teflon cuffs were secured to the pulmonary artery, pulmonary vein, and bronchus, and the graft was preserved on ice. Recipient mice were anesthetized, intubated, and mechanically ventilated before undergoing a left thoracotomy through the third and fourth intercostal space. Donor lungs were implanted using the cuff technique and secured with 10-0 Ethilon suture. The thoracotomy was closed using 6-0 suture.

The study included 26 mice: 12 donors, 12 transplant recipients, and two naïve controls. Each transplantation involved one donor and one recipient, with three recipients initially assigned to each experimental condition and collection time point. Donor and recipient animals were randomly selected from the appropriate strains. One day-28 isograft recipient was found dead by veterinary staff before the scheduled collection, with no documented preceding signs meeting humane-endpoint criteria. The cause of death was undetermined. Consequently, the final sample sizes were two day-28 isografts and three each for day-56 isografts, day-28 allografts, and day-56 allografts. Two naïve lungs served as baseline controls.

At scheduled experimental endpoints, surviving transplant recipients were euthanized by intraperitoneal administration of pentobarbital (100 mg/kg in 500 µL sterile saline) before lung harvesting for histology, flow cytometry, and TCRβ repertoire analysis. Donor animals (n = 12) underwent terminal procedures for lung procurement, and naïve controls (n = 2) were euthanized for baseline tissue collection. Death was confirmed using at least two criteria, including cessation of breathing and circulation, absence of corneal or withdrawal reflexes, or other established signs of death.

### Postoperative monitoring and humane endpoints

Following transplantation, recipients received 1 mL sterile saline subcutaneously, recovered on warming pads, and were administered buprenorphine (0.1 mg/kg, intraperitoneally) every 12 hours for postoperative analgesia. Animals were monitored daily for respiratory distress, changes in activity or appearance, and other signs of pain or postoperative complications.

The IACUC-approved protocol specified humane endpoints, including weight loss exceeding 20%, poor body condition, respiratory distress, and impaired mobility or responsiveness. Animals meeting these criteria were to undergo prompt veterinary evaluation and euthanasia when indicated. No animals were documented as requiring euthanasia after reaching humane-endpoint criteria. One recipient was found dead before the scheduled experimental endpoint, and the timing of any preceding deterioration could not be determined. Spontaneous death was not an intended experimental endpoint.

Research personnel received institutional training in animal care, handling, welfare, and relevant surgical procedures, including hands-on training appropriate to their responsibilities. Veterinary staff provided animal health oversight throughout the study.

### Lung histopathology

To assess graft injury, the left lungs were fixed overnight in 4% paraformaldehyde and processed for formalin-fixed paraffin embedding (FFPE) by the UCLA Translational Pathology Core Laboratory. Paraffin blocks were sectioned and stained with hematoxylin and eosin (H&E). Histologic images were acquired using a Zeiss Axioskop 2 Plus microscope and processed with AxioVision 4.2 software. Low-magnification (50x) images were used to evaluate overall tissue architecture, whereas high-magnification (200x) images were used to assess features of chronic inflammation and fibrosis.

### T-cell isolation

Naïve lungs, isografts, and allografts were mechanically dissociated through metal mesh and washed in MACS buffer containing 2% bovine serum albumin. CD90.2 cells were isolated with CD90.2 MicroBeads and LS columns (Miltenyi Biotec) according to the manufacturer’s protocol. When cell yield permitted, an aliquot was reserved for flow cytometry and the remaining cells were pelleted for TCRβ analysis.

### Flow cytometry

CD90.2-enriched cells were stained with fluorochrome-conjugated antibodies to CD45 (30-F11; BD 553081; RRID:AB_394611), CD3 (145-2C11; BD 553067; RRID:AB_394599), CD4 (RM4-5; BD 553046; RRID:AB_394582), and CD8 (53-6.7; BD 553035; RRID:AB_398527). Samples were acquired on a FACSCalibur cytometer and analyzed with CellQuest. CD4 and CD8 cells were reported as percentages of CD45 CD3 cells.

### Human CTOT study

CTOT-20 (NCT02631720, UCLA IRB 15-001574) prospectively enrolled 803 adults undergoing first lung transplantation at Duke University, UCLA, the University of Toronto, Johns Hopkins University, and Cleveland Clinic between December 22, 2015, through August 17, 2018 [8]. All participants provided written informed consent prior to study participation. BAL specimens were collected and processed as described previously [3].

### BAL collection and processing

During surveillance bronchoscopy, two 50–60 mL aliquots of isotonic saline were instilled into a subsegment of the right middle lobe or lingula [3]; investigators could adjust the volume to patient tolerance per ISHLT guidance [22]. Recovered fluid was pooled and divided for clinical and research use. Research BAL was kept on ice, processed within 6 h, filtered through sterile gauze, and centrifuged. BAL cell pellets (BAL-cp) were washed, preserved in RNAlater, and stored at −80 °C. Starting April 30^th^, 2018, gDNA was isolated from archived BAL-cp using the AllPrep DNA/RNA/miRNA Universal kit (Qiagen, Germany, Cat No. 80224) according to the manufacturer’s protocol. The derivative gDNA samples were placed back in storage for later research utilization.

### Patient and sample selection

After ethics approval at UCLA (UCLA IRB 19-0198) this CTOT ancillary study utilized a subset of nested CTOT-20 participants and their archived BAL-cp gDNA samples.

Selected participants were required to have at least 1 year of available follow-up and no documented respiratory infection during the first post-transplant year. Archived BAL-cp collected at approximately 1, 3, 6, 9, and 12 months post-transplant were considered. Samples required a gDNA concentration >1.0 ng/µL in a 95 µL elution for immunoSEQ library preparation, and participants were required to have at least two eligible samples. The final cohort comprised 23 recipients and 90 BAL-cp samples.

CLAD status was ascertained as described previously [8]. ACR grades were assigned to transbronchial biopsies obtained at the corresponding bronchoscopy using ISHLT criteria [7]. Recipients with any A1–A3 biopsy were classified as rejectors. Those without an A-grade event were classified as no-rejection recipients (labeled “Healthy” in the analysis). AX biopsies were not counted as events. For rejectors, the first A1–A3 observation was incident rejection. A0 observations before or after it were classified as pre-rejection or resolution, respectively. Later A1–A3 observations were classified as non-resolution if rejection persisted or recurrence if they followed an intervening A0 biopsy.

### TCRβ repertoire sequencing

gDNA was analyzed with the immunoSEQ Assay (Adaptive Biotechnologies) using multiplex PCR of TCRβ CDR3 rearrangements with bias-control methods [23]. Sequencing reads were demultiplexed, filtered, collapsed, and annotated with the immunoSEQ Analyzer to identify productive CDR3 amino acid clonotypes and their frequencies. Productive clonality was defined as 1 minus productive Shannon entropy normalized by the log -transformed number of unique productive rearrangements; values range from 0 (even) to 1 (oligoclonal) [13]. Maximum productive frequency was the frequency of the most abundant productive clonotype.

To describe assay input and repertoire sampling, DNA concentration and mass, template and rearrangement counts, productive fraction, assay-estimated cell measures, and productive clonality for all 90 samples from 23 recipients were summarized. Values are reported as sample-level medians (interquartile ranges).

### Murine repertoire overlap and shared-clonotype analyses

Day-28 and day-56 samples were pooled within the isograft (n = 5) and allograft (n = 6) groups. Abundance-weighted overlap was quantified with Morisita indexes exported from the immunoSEQ matrix. Each unordered mouse pair was included once; reciprocal entries and self-comparisons were excluded. Because mice contributed to multiple pairwise values, the group comparison was exploratory. For matched-size descriptive analyses, six five-of-six allograft leave-one-out (LOO) subsets, generated by leaving out one allograft at a time, were compared with all five isografts. Shared clonotypes required an identical productive CDR3 amino-acid sequence, matching V and J assignments, and productive frequency >0.001 wherever detected. Counts were calculated for clonotypes present in at least 2, 3, 4, or 5 mice. Shared clonotypes were ranked by aggregate productive frequency across samples, and the top 10 were retained. Allograft summaries were averaged across the six overlapping LOO subsets and treated as descriptive sensitivity analyses rather than independent comparisons.

### Month-6 surveillance-visit index analysis

Month 6 was selected a priori, before CLAD-outcome analyses, because it was a protocol-defined surveillance visit beyond the perioperative primary graft dysfunction window [24,25] and after recipient-derived graft T-cells begin acquiring tissue-resident phenotypes [26]. The exploratory index analysis included recipients with an eligible month-6 BAL-cp who were CLAD-free and under observation at collection (20 of 23 recipients; post-transplant days 137–223). The actual collection date defined index day 0. Productive clonality above the cohort median of 0.0743 was classified as high; values ≤0.0743 were classified as low.

Two CLAD endpoints were examined: Early-onset CLAD and CLAD at any timepoint. Early-onset CLAD, defined as CLAD within 2 years of transplantation [27]. Events diagnosed by post-transplant day 730 were counted; recipients without CLAD were administratively censored at day 730. Recipients were followed from the month-6 index BAL-cp until CLAD diagnosis or their last clinical follow-up.

### First-ACR index-event analysis

The exploratory first-rejection analysis was restricted to rejectors with an eligible incident-rejection BAL-cp. Index day 0 was the collection date of the earliest BAL-cp paired with an A1–A3 biopsy. Pre-rejection, recurrence, non-resolution, and resolution samples were ineligible as index samples, and recipients without an eligible index sample were excluded.

Eleven rejectors met these criteria. Productive clonality at the index ACR sample was classified using the month-6 threshold carried forward from the full cohort: high if >0.0743 and low if ≤0.0743. Follow-up began at the patient-specific index biopsy date and ended at CLAD diagnosis or the last available clinical follow-up.

### Statistical analysis

Analyses were performed using GraphPad Prism 11 and R 4.5.1. Two-group comparisons used two-sided unpaired Mann–Whitney U tests. For murine analyses, Cliff’s δ was also reported as a distribution-free effect size [28]. Longitudinal productive clonality was modeled on the raw scale with recipient-specific random intercepts using lme4 and lmerTest. For the exploratory CLAD analysis, visits were grouped into early (months 1 and 3) and late (months 6, 9, and 12) periods. The model included fixed effects for subsequent CLAD status, period (early vs. late), and their interaction. The CLAD-status-by-period interaction tested whether the early-to-late change in productive clonality differed between CLAD and CLAD-free recipients. The rejection-status model included fixed effects for rejection group and categorical month (1, 3, 6, 9, and 12). Mixed models were fitted by restricted maximum likelihood, and Satterthwaite tests were used for inference. CLAD-free survival was estimated by Kaplan–Meier analysis, and groups were compared using two-sided log-rank tests with survival 3.8-3. Data are shown as mean ± SEM unless stated otherwise. P values were two-sided and unadjusted for multiple comparisons; P < 0.05 was considered significant.

### Ethics statement

All animal procedures were approved by the University of California, Los Angeles (UCLA) Institutional Animal Care and Use Committee (IACUC; protocol ARC-2010-057) and conducted in accordance with the National Institutes of Health Guide for the Care and Use of Laboratory Animals and institutional animal welfare standards. The approved protocol included appropriate anesthesia, postoperative analgesia, daily monitoring, and predefined humane endpoints to minimize pain and distress. Spontaneous mortality was not an intended experimental endpoint, and any unexpected deaths were documented and reported as part of the study outcomes. The human CTOT-20 study was approved by the institutional review boards of the participating study centers, including the University of California, Los Angeles Institutional Review Board (UCLA IRB 15-001574), and all participants provided written informed consent prior to study participation. The present ancillary study involving archived bronchoalveolar lavage samples and associated clinical data was approved by the UCLA Institutional Review Board (IRB 19-0198). All human-subject research was conducted in accordance with applicable institutional requirements and the principles of the Declaration of Helsinki.

## Results

### Lung allografts exhibit chronic inflammatory remodeling and altered T-cell composition

To establish the experimental context for TCR repertoire analysis, we used a murine orthotopic left lung transplant model comparing C57BL/6 lungs transplanted to C57BL/6 recipients (isografts) with BALB/c lungs transplanted to C57BL/6 recipients (allografts). Days 28 and 56 were selected to evaluate established graft responses beyond the peri-transplant period, when ischemia–reperfusion injury may predominate [24]. We hypothesized that isografts and naïve lungs would be virtually indistinguishable at these time points, whereas allografts would show marked injury. Naïve control lungs were collected at baseline, and transplanted lungs were harvested for histologic evaluation, flow-cytometric characterization, and TCRβ CDR3 sequencing (Fig 1A).

**Fig 1.**
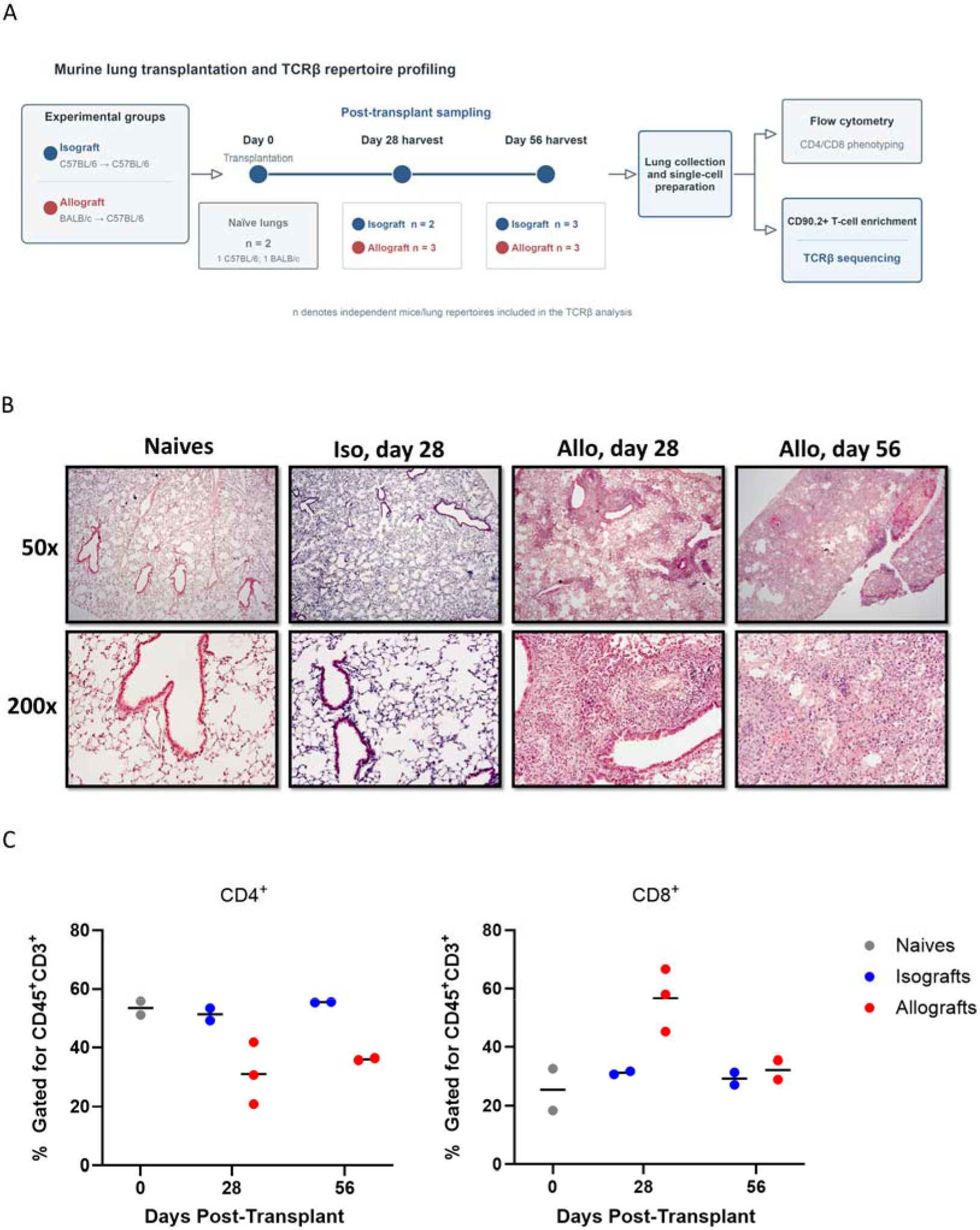
Murine orthotopic lung transplantation induces progressive allograft injury and enables characterization of graft-infiltrating T-cells. (A) Experimental design comparing allografts (Allo), isografts (Iso), and naïve lungs at the indicated post-transplant time points. (B) Representative hematoxylin and eosin (H&E)-stained sections of naïve lungs and isografts and allografts collected at days 28 and 56 after transplantation. (C) Flow cytometric characterization of lung-isolated CD90.2□-purified T-cells. CD4□ (left) and CD8□ (right) T-cells, expressed as percentages of CD45^⁺^CD3^⁺^ cells, in CD90.2^⁺^-enriched preparations from naïve, isograft, and allograft lungs used for TCR repertoire sequencing. Data are presented as mean ± SEM, with individual animals represented by dots.

Histopathologic evaluation included naïve lungs (1 BALB/c and 1 C57BL/6), isografts collected at days 28 (n = 2) and 56 (n = 3), and allografts collected at days 28 (n = 3) and 56 (n = 3). Representative H&E-stained sections of the naïve left lungs and left isografts at days 28 and 56 demonstrated preserved arteries and veins without reactive endothelial cells or infiltrating leukocytes. The airways were normal without epithelial hyperplasia, injury or inflammation. The alveolar architecture was preserved, with normal-appearing pneumocytes and an unremarkable interstitium without stromal expansion or inflammatory cell infiltration (Fig 1B). In contrast, day-28 allografts exhibited perivascular and peribronchial mononuclear cell infiltration. There is associated endothelial cell and airway epithelial cell reactivity and damage. The interstitium demonstrated inflammation with pneumocyte reactivity and damage. By day 56, allografts displayed continued inflammation with the addition of stomal cells surrounding the vessels, airways and interstitium causing architectural distortion (Fig 1B). These findings demonstrated allograft injury and structural remodeling in the experimental setting used for subsequent repertoire analysis.

We confirmed the presence of both CD4□ and CD8□T-cell populations in the preparations used for subsequent TCR repertoire analysis (Fig 1C). Although variation in the relative proportions of CD4□ and CD8□ cells was observed among experimental groups and time points, these analyses were performed to characterize the T-cell composition of the purified preparations rather than to assess transplantation-associated changes in T-cell subset distribution. Together, these data confirmed that the lung-derived CD90.2□-purified preparations contained both major conventional T-cell subsets prior to TCR repertoire sequencing.

### Lung allografts exhibit increased intragraft TCR clonality and dominant-clonotype frequency

We next sought to determine whether graft-dependent changes in T-cell populations were accompanied by alterations in T-cell repertoire composition and clonal expansion. Productive TCRβ repertoires were analyzed in CD90.2-enriched T-cells from naïve lungs, isografts collected at days 28 (n = 2) and 56 (n = 3), and allografts collected at days 28 (n = 3) and 56 (n = 3). Productive clonality ranges from 0 to 1, with higher values indicating a more uneven repertoire dominated by fewer clonotypes [13]. Naïve lungs served as baseline references, whereas inferential comparisons focused on isografts and allografts to control for transplantation-associated effects. Day 28 was selected to minimize contributions from perioperative and ischemia-reperfusion injury. Naïve and isograft repertoires appeared comparable, although the small sample sizes precluded definitive statistical assessment.

For an overall comparison, samples collected at days 28 and 56 were pooled within the isograft (n = 5) and allograft (n = 6) groups, with individual-time point analyses performed to evaluate the consistency of the observed differences. Productive clonality was significantly higher in allografts than in isografts (Cliff’s δ = 0.93, p = 0.0087; Fig 2A), demonstrating greater repertoire skewing. For context, δ = 0.93 represents near-complete rank separation, while δ = 1.00 would indicate that every observed value in the allograft group exceeded every corresponding value in the isograft group. When the two time points were analyzed separately, productive clonality remained higher in allografts than in isografts at both days 28 and 56 (Fig 2B). The observed values showed complete rank separation between groups at both time points (day 28: Cliff’s δ = 1.00, p = 0.20; day 56: Cliff’s δ = 1.00, p = 0.10).

**Fig 2.**
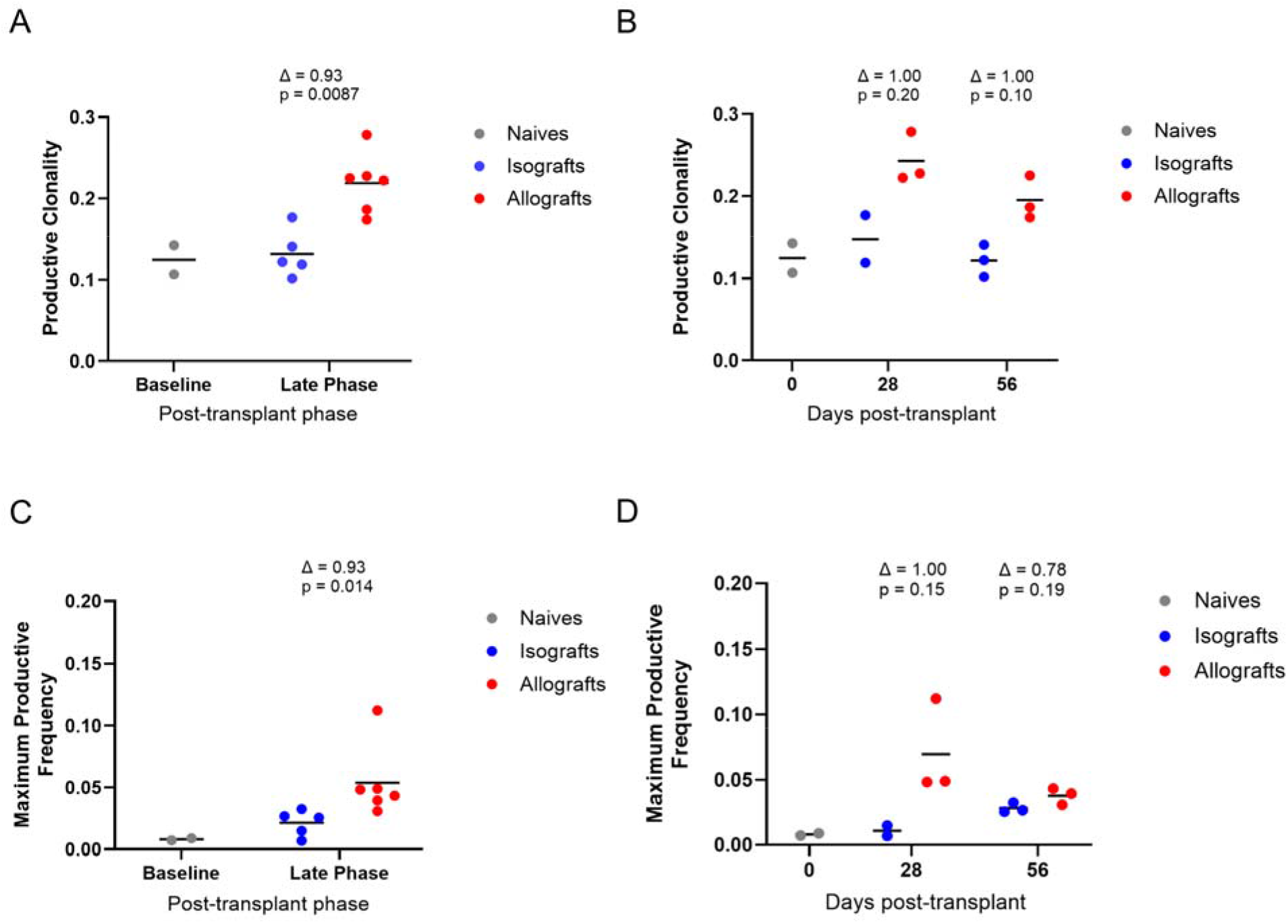
Lung allografts exhibit persistent T-cell clonal expansion and repertoire remodeling. (A) Productive clonality in pooled day-28 and day-56 isografts and allografts. Day 28 and day 56 were combined to represent a biologically similar phase of established alloimmune responses characterized by expansion of T-cells during allograft rejection (Cliff’s δ = 0.93, p = 0.0087). (B) Productive clonality at each time point (day 28: δ = 1.00, p = 0.20; day 56: δ = 1.00, p = 0.10). (C) Maximum productive frequency in pooled day-28 and day-56 samples (δ = 0.93, p = 0.014). (D) Maximum productive frequency at each time point (day 28: δ = 1.00, p = 0.15; day 56: δ = 0.78, p = 0.19). Isograft/allograft sample sizes were 2/3 at day 28, 3/3 at day 56, and 5/6 in pooled analyses.

Maximum productive frequency, representing the relative abundance of the most frequent productive clonotype in each repertoire, was also significantly higher in combined allografts (Cliff’s δ = 0.93, p = 0.014; Fig 2C). This finding was consistent with increased representation of dominant clonotypes within the allograft repertoire. When the two time points were analyzed separately, maximum productive frequency showed the same directional pattern, with higher values in allografts at day 28 (Cliff’s δ = 1.00, p = 0.15) and day 56 (Cliff’s δ = 0.78, p = 0.19) (Fig 2D).

Together, these findings indicate that lung allografts exhibit increased intragraft T-cell repertoire skewing and greater representation of dominant clonotypes. Although the antigen specificity of the expanded clonotypes was not determined, the observed pattern is consistent with selective clonal expansion within the allograft environment.

### Lung allografts exhibit greater repertoire overlap and enrichment of recurrent clonotypes

We next examined whether this repertoire remodeling was accompanied by increased repertoire overlap and recurrence of shared clonotypes across independent recipients. We compared TCRβ repertoire overlap among isografts and allografts collected at days 28 and 56. This inbred murine model allowed repertoire similarity to be examined among genetically matched C57BL/6 recipients exposed to lungs from the same BALB/c donor strain. Because lung harvesting was terminal, overlap was calculated across independent recipients rather than longitudinally within animals. Pairwise Morisita overlap was greater among allografts than among isografts (isograft n = 5, allograft n = 6; Mann–Whitney p < 0.0001; Fig 3A). Each unordered mouse pair was counted once, with reciprocal matrix entries excluded. This exploratory comparison supports greater abundance-weighted repertoire overlap among allografts compared to isografts.

**Fig 3.**
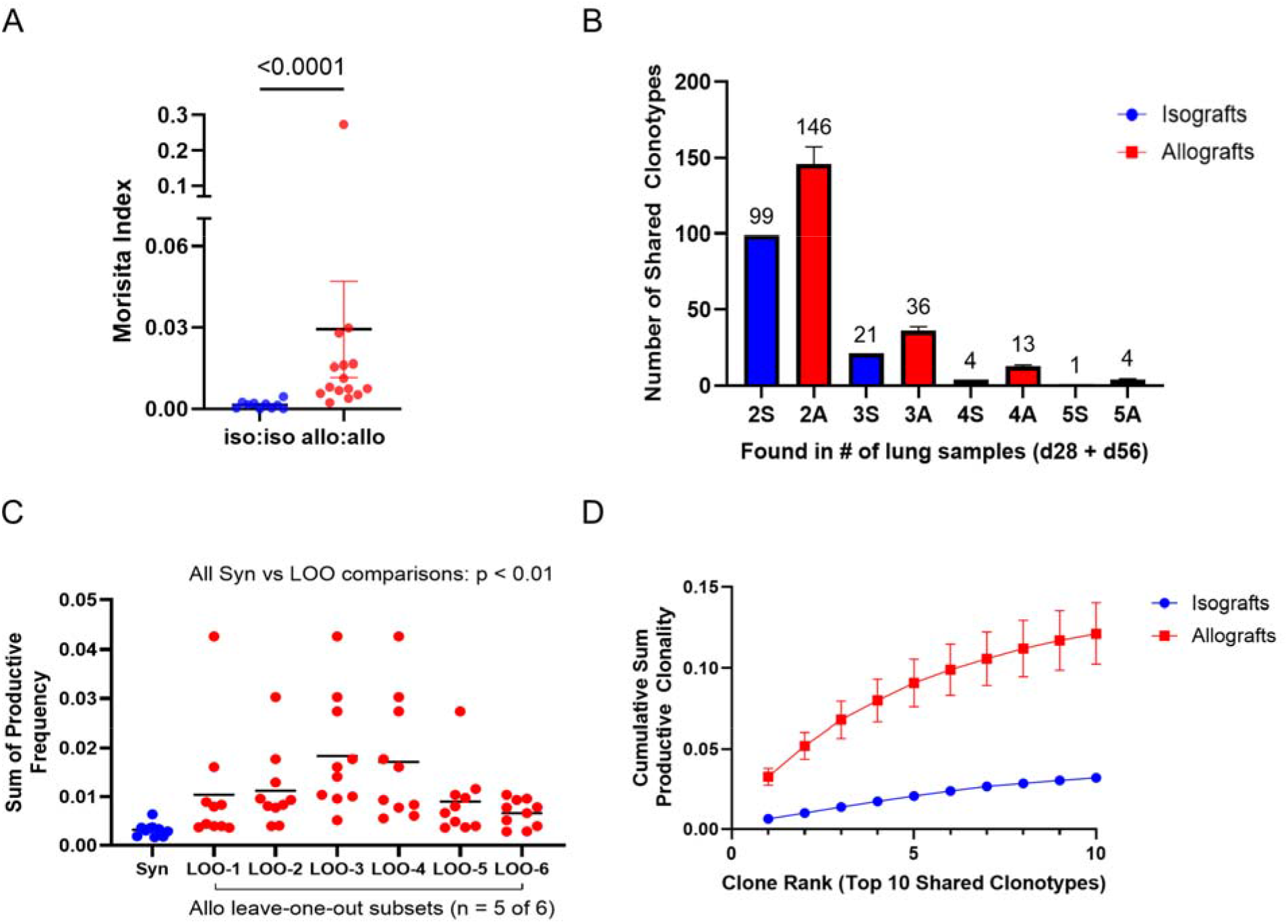
Lung allografts exhibit greater repertoire overlap and enrichment of shared T-cell clonotypes. (A) Pairwise Morisita overlap among pooled day-28 and day-56 isografts (n = 5) and allografts (n = 6). Each unordered mouse pair was plotted once, with reciprocal entries excluded (p < 0.0001, two-sided Mann–Whitney U test). Because mice contributed to multiple pairs, this comparison was exploratory. (B) The number of shared clonotypes detected in any 2, 3, 4, or 5 pooled lung samples. Shared clonotypes are defined by having identical productive CDR3 amino-acid sequences, matching V and J assignments, and productive frequency >0.001 in all detected samples. To account for unequal group sizes (five isografts vs six allografts), each allograft was sequentially excluded to generate six matched leave-one-out (LOO) subsets (LOO-1–LOO-6; n = 5 each). Isograft (I) values encompass all five isografts; allograft (A) values are means across all six LOO allograft subsets. Whiskers show variation across the six overlapping subsets. (C) Aggregate productive frequency of the top 10 shared clonotypes. The isograft curve includes all five mice; the allograft curve is the mean across the six LOO subsets. Frequencies were summed across samples in which each clonotype was detected; points represent clonotypes and horizontal lines indicate means. Exploratory analysis comparing 10 clonotype aggregate productive frequency of isografts differed significantly from all six LOO subsets (two-sided unpaired Mann– Whitney U tests: LOO-1, p = 0.0007; LOO-2–4, p < 0.00001; LOO-5, p = 0.0005; LOO-6, p = 0.0089). (D) Cumulative aggregate productive frequency of the top 10 shared clonotypes. The isograft curve includes all five mice; the allograft curve is the mean across the six LOO subsets, with whiskers showing variation. Panels B–D are descriptive sensitivity analyses because the allograft subsets overlap and are not independent biological groups.

We next quantified shared clonotypes in matched groups of five recipient mice using the prespecified sequence, V/J-assignment, and productive-frequency criteria of intragraft T-cells. In the five isografts, 99, 21, 4, and 1 clonotypes were shared in any 2-, 3-, 4-, and 5-pooled lung samples, respectively. Across the six leave-one-out (LOO) allograft subsets the corresponding mean shared clonotypes were 146, 36, 13, and 4 clonotypes, respectively, shared in any 2, 3, 4, and 5 pooled lung samples (Fig 3B). The allograft subsets, excluding a different allograft, were created with the intent to generate matched-size results to the isograft group as an exploratory descriptive sensitivity analysis. All subsets showed the trend of increased recurrent clones among allografts compared to isografts.

To assess the abundance of the dominant recurrent populations, we ranked shared clonotypes within each group or subset by aggregate productive frequency and examined the top 10. The summed productive frequencies of these clonotypes were greater in all six leave-one-out allograft subsets than in the five isografts (Fig 3C). Rank-based cumulative analysis likewise showed a greater cumulative aggregate productive frequency across the top 10 shared clonotypes in the matched allograft subsets than in isografts (Fig 3D). This descriptive pattern indicated that a limited number of recurrent T-cell populations accounted for a larger aggregate share of the productive repertoire under allograft conditions.

Together, the exploratory pairwise Morisita analysis showed greater abundance-weighted repertoire overlap among allografts, as compared to isografts, with each unordered pair included once. The matched-size descriptive analyses quantified the recurrent clonotypes among respective subsets and suggested greater representation of dominant recurrent T-cell populations in allografts. This pattern is consistent with convergent or public T-cell responses under shared antigenic pressures in the allograft environment [29–31].

### BAL-cp productive clonality at month 6 differs by CLAD outcome and stratifies survival free from early onset CLAD

To investigate whether the repertoire remodeling observed in the murine model was also evident in human lung transplantation, we analyzed longitudinal BAL samples from 23 lung transplant recipients undergoing surveillance bronchoscopy. Seventeen recipients remained CLAD-free, whereas 6 developed CLAD, including 4 who developed early-onset CLAD defined as within 24 months of transplantation (Fig 4A). Additional patient characteristics and sample-level biopsy findings are provided in S2 Table.

**Fig 4.**
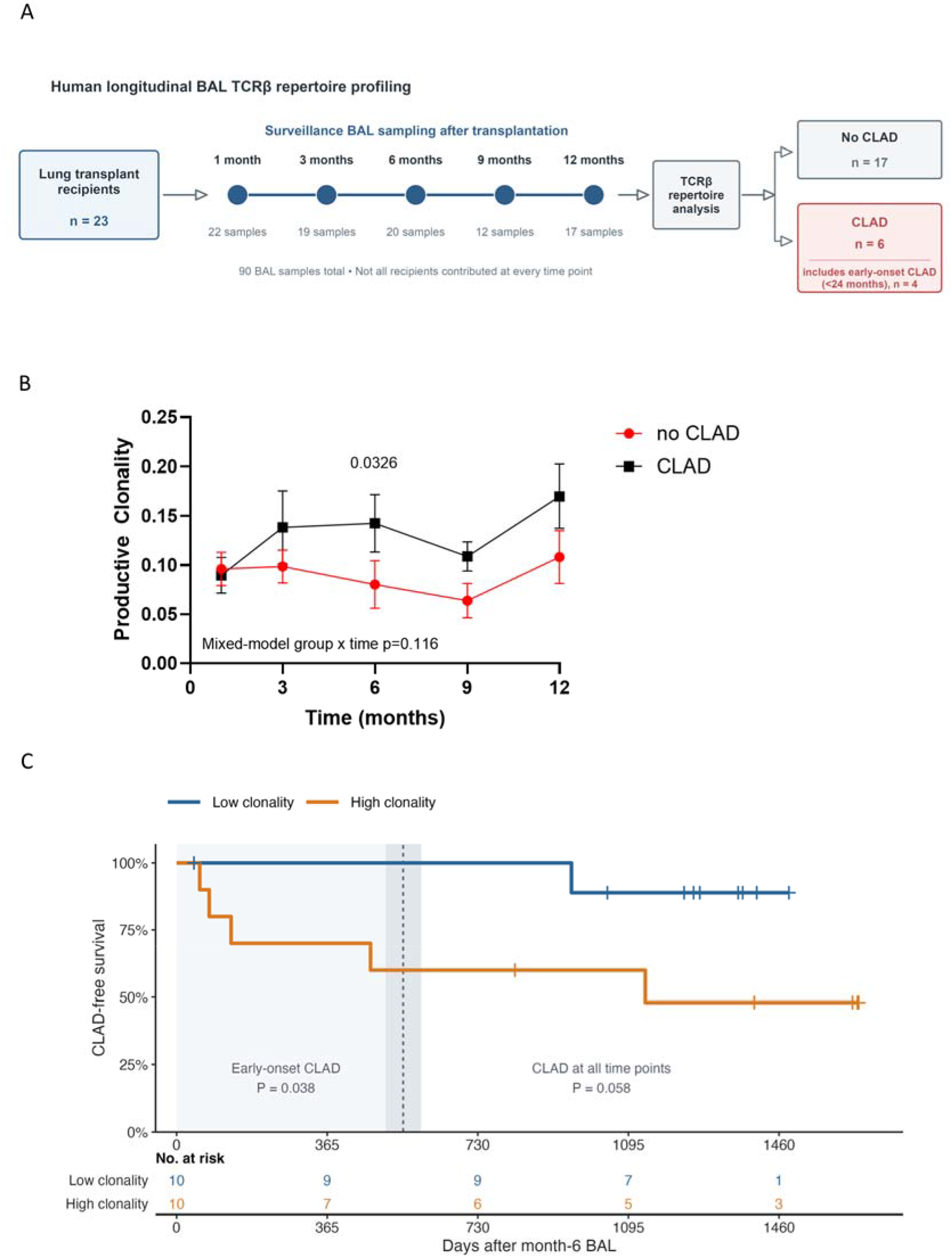
Longitudinal BAL-cp productive clonality differs by CLAD outcome at month 6 and stratifies early-onset CLAD-free survival. (A) BAL-cp collected approximately 1, 3, 6, 9, and 12 months after lung transplantation underwent TCRβ sequencing. Recipients were classified as CLAD-free or CLAD, including early-onset CLAD (<24 months). Missing visits reflected limited BAL-cp cellularity, insufficient DNA yield, or clinical sampling. (B) Mean ± SEM productive clonality by subsequent CLAD status. At the prespecified month-6 visit, productive clonality was higher in recipients who developed CLAD (p = 0.0326, two-sided unpaired Mann–Whitney U test). An exploratory raw-scale linear mixed-effects model included CLAD status, period (early: months 1 and 3; late: months 6, 9, and 12), their interaction, and a recipient-specific random intercept; the group × period interaction was not significant (p = 0.116). (C) Kaplan–Meier estimates among 20 recipients with eligible month-6 BAL-cp who remained CLAD-free at that visit. The collection date (post-transplant days 137–223) defined the index day 0. Productive clonality was classified as high (>0.0743) or low (≤0.0743). The shaded region denotes the early-onset window: events occurred in 0 of 10 low and 4 of 10 high recipients (p = 0.038). Across all follow-up time, events occurred in 1 of 10 low and 5 of 10 high recipients (p = 0.058). Survival comparisons used two-sided log-rank tests. Tick marks denote censoring; tables show numbers at risk.

We compared productive clonality longitudinally in recipients classified according to CLAD status during follow-up. Assay metrics for all 90 BAL-cp samples and the month-6 surveillance samples are shown in S1 Table. At month 6, median productive template counts were 2,620 (interquartile range, 2,281–4,073) among recipients who subsequently developed CLAD and 1,309 (442–2,926) among those who remained CLAD-free. Month-6 surveillance visit BAL cell-pellet productive clonality was higher among recipients who subsequently developed CLAD (n = 6; median, 0.1424) than among those who remained CLAD-free (n = 14; median, 0.0486; Hodges–Lehmann pairwise difference, 0.0643; 95% CI, 0.0053–0.1634; two-sided unpaired Mann–Whitney U test, p = 0.0326; Fig 4B). In an exploratory mixed-effects analysis, visits were grouped into early (months 1 and 3) and late (months 6, 9, and 12) periods. Model-estimated productive clonality was 0.102 during the early period and 0.144 during the late period among recipients who developed CLAD, compared with 0.098 and 0.088, respectively, among CLAD-free recipients. The estimated between-group difference in early-to-late change was 0.052 (95% CI, −0.013 to 0.116), but the CLAD-status-by-period interaction was not statistically significant (p = 0.116). Thus, the data showed a directional pattern of divergence but did not establish that productive clonality changed differently according to subsequent CLAD status.

Using the cohort-derived median cutoff of 0.0743, early-onset CLAD occurred in 4 of 10 high-clonality recipients and none of 10 low-clonality recipients (log-rank p = 0.038). Across all follow-up time, CLAD occurred in 5 of 10 and 1 of 10 recipients, respectively (p = 0.058). This cohort-derived threshold analysis was exploratory.

### BAL-cp productive clonality is associated with CLAD-free survival among recipients with ACR

We next examined whether BAL-cp productive clonality was associated with biopsy-defined ACR. Across 90 paired BAL-biopsy observations, 68 biopsies were classified as A0, 16 as A1–A3, and 6 as AX. At the recipient level, 12 of 23 recipients experienced at least one A1–A3 rejection event and were classified as rejectors, whereas 11 had no biopsy-defined A-grade rejection event and were classified as healthy/no-rejection recipients (S3 Table).

Productive clonality measurements of healthy patients and patients with ACR showed no consistent separation when plotted longitudinally across surveillance months 1, 3, 6, 9, and 12 months (Fig 5A). These findings suggest that isolated cross-sectional measurements incompletely capture the heterogeneous and dynamic nature of ACR following lung transplantation.

**Fig 5.**
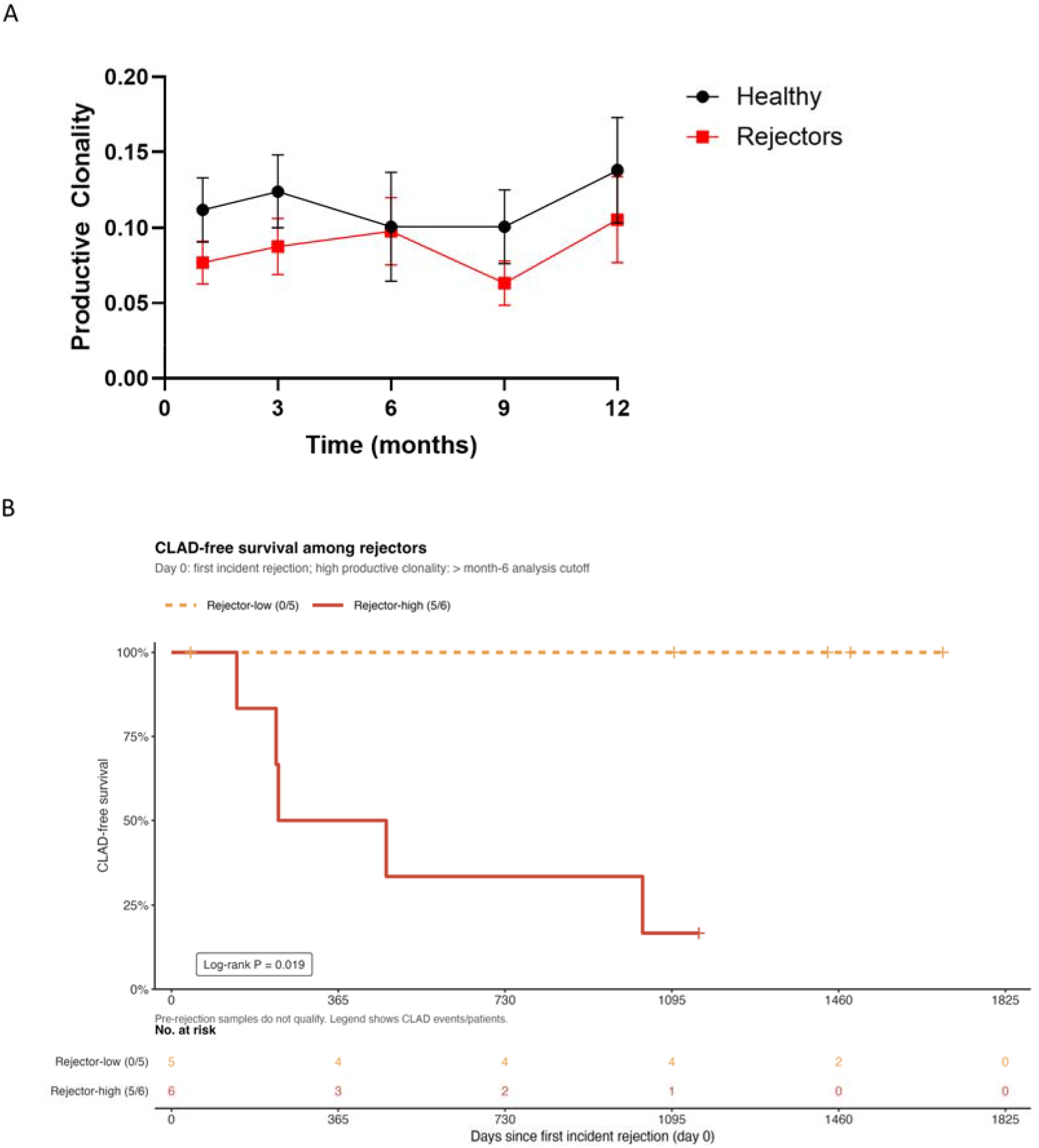
Patient-specific rejection status and productive clonality joint stratification is associated with CLAD-free survival. (A) Productive clonality at months 1, 3, 6, 9, and 12 in no-rejection recipients (labeled Healthy) and patients with ACR (rejectors). Groups overlapped, with no significant difference at any individual visit. (B) CLAD-free survival among 11 rejectors with biopsy-defined A1–A3 rejection and an eligible incident-rejection BAL-cp. Index day 0 was the earliest incident-rejection BAL collection; pre-rejection, recurrence, non-resolution, and resolution samples were excluded. The month-6 cutoff from Fig 4C was carried forward: high, >0.0743; low, ≤0.0743. CLAD occurred in 0 of 5 low and 5 of 6 high recipients (p = 0.019, two-sided log-rank test). Tick marks denote censoring; the table shows numbers at risk. Source data are provided in the Source Data file.

Because productive clonality alone showed limited separation by rejection status, we used a patient-specific index-event analysis to examine whether the ACR status and productive clonality were associated with CLAD-free survival. For each rejector, productive clonality was measured at the first eligible incident-rejection BAL specimen. Among the 11 eligible rejectors, productive clonality status was assigned into high (n = 6; 5 CLAD events) and low (n = 5; 0 CLAD events) using the total cohort cutoff of 0.0743. CLAD-free survival differed between the high- and low-clonality groups significantly (log-rank p = 0.019; Fig 5B).

Thus, productive clonality alone showed limited separation by recipient-level rejection status. At patient-specific index events, however, CLAD events were concentrated among rejectors with high productive clonality. This exploratory analysis used cohort-derived thresholds and different index definitions for rejectors and healthy/no-rejection recipients.

## Discussion

We combined a controlled murine orthotopic lung transplant model with longitudinal BAL-cp TCRβ profiling in human recipients to investigate graft-proximal TCRβ repertoire remodeling and its relationship to acute and chronic allograft injury. Murine allografts showed greater repertoire skewing, dominant-clonotype expansion, and abundance-weighted overlap than isografts, although the pairwise overlap analysis was exploratory. In humans, productive clonality showed limited separation by biopsy-defined ACR but was higher at month 6 among recipients who later developed CLAD. CLAD events were also concentrated among rejectors with high productive clonality at first incident rejection. Together, these findings support the potential of BAL-cp TCR repertoire remodeling as a longitudinal correlate of chronic allograft injury that may complement histologic rejection assessment.

The murine transplant model provides experimental evidence that the allograft environment can reshape the local T-cell repertoire. Compared with isograft controls, allografts showed increased productive clonality and higher maximum productive frequency, indicating preferential expansion of dominant clonotypes. In addition, pairwise Morisita overlap was greater among allografts, and matched-size descriptive analyses showed more shared clonotypes and greater representation of the top shared clonotypes in allografts than in isografts. Although antigen specificity was not directly assessed, the enrichment of recurrent dominant clonotypes is consistent with antigen-driven or public T-cell responses arising in the allograft environment [29–31]. The human transplant patient cohort extended these observations into a clinically relevant setting, but with important differences from the controlled murine model. Productive clonality did not robustly distinguish healthy/no-rejection recipients from recipients with rejection when analyzed across isolated surveillance samples. This negative finding is biologically informative. ACR is episodic, spatially heterogeneous, and diagnosed by biopsy, which samples only a limited portion of the allograft and is subject to interpretive variability [7,9,10]. In addition, BAL-cp repertoires in human recipients may be influenced by infection and other inflammatory or non-alloimmune exposures [17,30]. Therefore, a single productive clonality measurement may be insufficient to identify biopsy-defined ACR at a given surveillance time point. In contrast, productive clonality showed a stronger relationship with future CLAD status. Month 6 was selected a priori, before inspection of productive-clonality differences by CLAD status or survival outcomes, because it was a protocol-defined surveillance visit beyond the immediate perioperative primary graft dysfunction window [24,25] and is a time when recipient-derived graft T-cells progressively acquire tissue-resident phenotypes over the first post-transplant months [26]. BAL-cp TCR repertoires at the month-6 time point demonstrated higher productive clonality among recipients who went on to develop early-onset CLAD, supporting our hypothesized index-visit selection. The absence of a significant group-by-time interaction indicates that this observation should be interpreted as a between-group difference at month 6, not as evidence of a formal temporal change point or of different overall trajectories.

As shown in Fig 4B, productive clonality did not visually separate recipients who developed CLAD from those who remained CLAD-free at the month-1 and month-3 visits. This early overlap may have attenuated evidence for divergent longitudinal trajectories and thereby contributed to the nonsignificant group-by-time interaction. One plausible biological explanation is that BAL-cp T-cell repertoires during the early post-transplant period reflect several concurrent immune processes rather than subsequent CLAD risk alone. Acute rejection is common during the first three months post-transplantation [32]. Pathogen-specific clones can expand in the graft during infection [17], and ACR can produce oligoclonal expansion of recipient-derived cytotoxic T-cells [18]. These competing immune signals could increase within-group heterogeneity and obscure an emerging CLAD-associated repertoire pattern. This interpretation remains speculative because infection and acute rejection were not modeled as time-varying covariates in the present analysis.

Infection and other non-alloimmune exposures remain possible beyond six months [33] and may influence BAL-cp repertoire measurements [17,30]. Greater productive clonality is biologically compatible with cumulative antigen exposure and expansion of dominant T-cell clones, although the donor specificity of these expanded populations was not established. These findings are consistent with prior BAL-based studies showing that graft-proximal immune and molecular features can precede or associate with lung allograft injury or dysfunction [3–5,20,34,35].

An important implication of these findings is that BAL-cp TCR repertoire profiling may complement conventional biopsy-based assessment. Productive clonality showed limited separation between healthy/no-rejection recipients and rejectors, indicating that it is unlikely to function as a standalone diagnostic marker of ACR. In the patient-specific index-event analysis, however, CLAD events occurred in 5 of 6 (83%) rejectors with high productive clonality measured at first incident rejection and in none of the 5 rejectors (0%) with low clonality. Productive clonality may therefore capture a dimension of graft-proximal immune remodeling that is not represented by histologic classification alone.

This study also supports the value of sampling the lung allograft compartment directly. Much transplant repertoire work has focused on peripheral blood, but circulating T-cell repertoires may not fully represent immune activity within the graft [13,36]. BAL provides a clinically accessible graft-proximal sample during routine surveillance bronchoscopy and may therefore be useful for longitudinal immune monitoring in lung transplantation [3–5,17,20,22,34,35]. The present findings suggest that longitudinal BAL-cp repertoire features, particularly measures of repertoire skewing and dominant clone expansion, may help identify immune states associated with adverse long-term allograft outcomes. Several limitations should be acknowledged. The modest murine and human sample sizes limited statistical power and precluded robust multivariable adjustment. The human cohort was observational and clinically heterogeneous, including variation in immunosuppression, infection history, sampling time, and other transplant-related exposures. BAL is graft-proximal but does not represent the entire allograft, and BAL-cp cellular composition may vary between samples. Productive clonality is not specific to alloimmunity and may also reflect infection, inflammation, or other antigenic stimuli [17,30]. Because donor and antigen specificity were not directly tested, increased repertoire overlap and shared clonotypes are consistent with, but do not establish, convergent alloimmune responses; public TCRs may also arise through convergent recombination or recurrent responses to common antigens [29–31]. The CLAD analyses were likewise exploratory, with six total CLAD events and four early-onset events, and the cohort-derived median threshold requires validation in an independent cohort.

The Fig 5B analysis has additional design constraints. It was restricted to the 11 rejectors with an eligible incident-rejection BAL sample. This patient-specific index aligns repertoire measurement with a clinically relevant rejection event but introduces heterogeneity in post-transplant timing and excludes recipients without an eligible index sample. The high- and low-clonality groups were small (n = 6 and n = 5, respectively), with five CLAD events in the high-clonality group and none in the low-clonality group. These sparse data precluded stable multivariable or interaction modeling. Moreover, the productive-clonality cutoff of 0.0743 was derived from the same overall cohort in the separate month-6 analysis, so the observed survival separation may be optimistic. The result should therefore be interpreted as event enrichment within an exploratory rejector subgroup, not as evidence that productive clonality modifies the effect of rejection, a validated prognostic threshold, or a causal relationship. Larger independent cohorts with prespecified thresholds, harmonized sampling windows, adjustment for infection and treatment exposures, single-cell TCR sequencing, cellular phenotyping, and donor-reactivity assays are needed to validate these findings.

In summary, our findings indicate that lung allografts exhibit local T-cell repertoire remodeling characterized by increased clonality, dominant-clonotype expansion, and repertoire convergence in a controlled pre-clinical murine model. In human lung transplant recipients, BAL-cp productive clonality did not robustly associate with biopsy-defined ACR but was higher at the month-6 visit among recipients who developed CLAD and was associated with early-onset CLAD in the month-6 surveillance-visit index analysis. In the patient-specific index-event analysis, productive clonality measured at first incident rejection identified an event-enriched rejector subgroup. These results suggest that BAL-cp TCR repertoire profiling may complement biopsy-based assessment as a longitudinal measure of chronic allograft immune remodeling and CLAD risk rather than serve as a standalone cross-sectional diagnostic test for ACR.

## Acknowledgments

The authors thank Ying Ying Xie for technical assistance with the murine lung transplantation studies. We gratefully acknowledge the contributions of the many individuals who helped make CTOT-20 possible, including the CTOT-20 site PI’s John Reynolds, Pali D. Shah, Lianne G. Singer, and Marie Budev, the contributions of the many individuals at the data coordinating center, Rho, and the individuals at the DCRI, including Megan Neely. The authors offer their sincere gratitude to the research study coordinators at each of the enrolling sites for their efforts and to the many physicians, nurses, and others who comprise the lung transplant clinical care teams at each CTOT-20 enrolling center, in addition to all the lung transplant recipients who participated in this study, without whom this research would not be possible. We thank the immunoSEQ technical support team at Adaptive Biotechnologies for technical guidance, feedback, and recommendations regarding TCR repertoire data analysis. We also thank the Saul and Joyce Brandman Foundation Center for Lung Health at UCLA for its support of this work.

## Supporting information captions

**S1 Table. BAL cell-pellet TCR**β **assay metrics across all 90 first-year samples and at the month-6 surveillance visit.** A. All follow-up BAL samples. B. Month-6 surveillance-visit BAL samples. Values are at the sample level. Because multiple BAL samples were collected from individual recipients, no p-values are reported for this descriptive sample-level table. **Notes.** Values are median (interquartile range). Panel A includes repeated samples from 23 recipients (17 without and 6 with CLAD during follow-up); two samples from the CLAD group were collected on or after CLAD diagnosis. Panel B contains one pre-CLAD sample per recipient; its CLAD grouping describes subsequent clinical status. Nucleated-cell and T-cell counts and T-cell fraction are assay estimates, not independent flow-cytometry measurements. Template counts are shown as the quantitative input measure.

**S2 Table. Patient-level characteristics of the human BAL TCR cohort stratified by CLAD status.** Values are patient-level unless otherwise specified. **Notes.** ATG, anti-thymocyte globulin; CLAD, chronic lung allograft dysfunction; CNI, calcineurin inhibitor; DSA, donor-specific antibody; IQR, interquartile range; MMF, mycophenolate mofetil; PGD, primary graft dysfunction. P values compare No CLAD versus CLAD groups and are exploratory: exact Mann–Whitney tests were used for continuous variables and Fisher’s exact tests for binary categorical variables. Multilevel descriptive categories and outcome-defining variables were not statistically tested. Patient-level ACR reflects any biopsy A-grade rejection event (A1–A3) in the available surveillance dataset; A0 and AX samples were not counted as A-grade rejection events.

**S3 Table. Sample-level biopsy characteristics of BAL samples used for human TCR analysis.** Values are at the sample level. Because multiple BAL samples were collected from individual recipients, no p-values are reported for this descriptive sample-level table. **Notes.** A-grade rejection scores are reported from transbronchial biopsy results paired with BAL surveillance samples. A1–A3 indicate biopsy-defined ACR grades represented in this dataset; A0 indicates no ACR; AX indicates ungradable biopsy. B-grade values summarize airway inflammation grading where available. Sample categories reflect the analysis labels in the source dataset and are provided to support interpretation of later ACR analyses.

